# DreaM: A Computational Pipeline for Enhanced Short-Read Sequence Analysis in Repetitive Genomic Regions

**DOI:** 10.1101/2024.11.12.623194

**Authors:** Santosh Kumar, Fumiko Esashi

## Abstract

Mapping short sequencing reads to repetitive genomic regions, such as centromeres, presents significant challenges, primarily due to PCR duplicates, which can be erroneously mapped multiple times within these regions. Conventional bioinformatics pipelines often overlook this issue, potentially leading to misinterpretation as signal enrichment. To address this, we developed **DreaM** (Deduplication of Reads for Enhanced and Accurate Mapping), a computational pipeline that prioritises the preprocessing of raw sequencing data. DreaM firstly identifies and removes PCR duplicates, which is followed by read trimming to reduce noise from multiply mapped reads. When applied to ChIP-Seq and CUT&RUN datasets targeting CENP-A, a key marker of centromeres, DreaM demonstrated improved peak detection within centromeres. Overall, DreaM provides a robust solution for enhancing the analysis of DNA-protein binding sites in repetitive genomic regions using short-read sequencing.

## Introduction

Next-generation sequencing-based DNA-protein interaction studies, such as ChIP-Seq (Chromatin Immunoprecipitation Sequencing)^1^, CUT&RUN (Cleavage Under Targets and Release Using Nuclease)^2,3^, CUT&TAG (Cleavage Under Targets and Tagmentation)^4^, and DamID (DNA adenine methyltransferase identification)^5^, are widely conducted to provide a genome-wide landscape of protein-DNA associations (**Fig.1A**).

**Figure.1:**
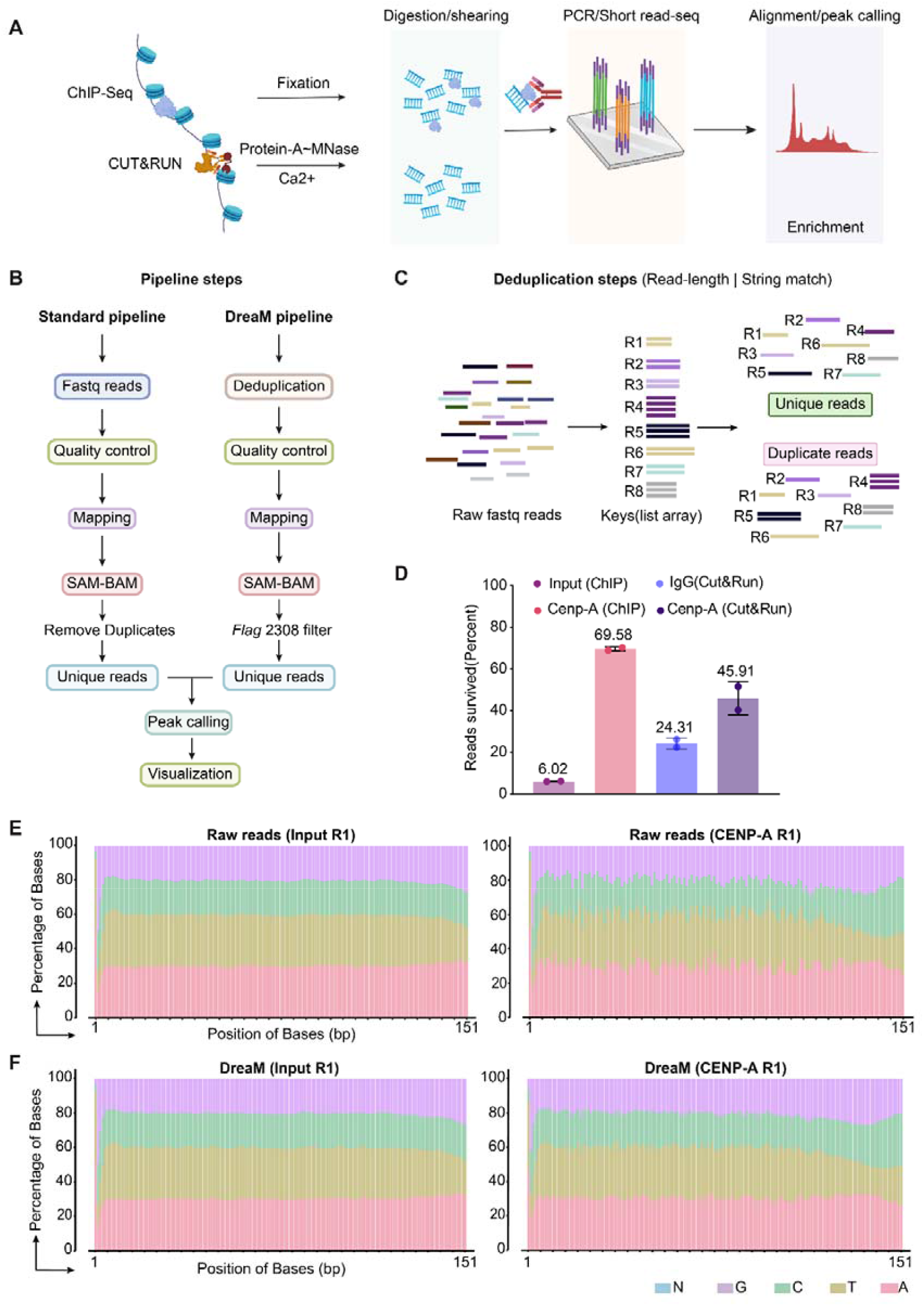
Diagrams encompassing **(A)** steps for major DNA-protein interaction studies using short-read sequencing, **(B)** Conventional pipeline used to analyze the data and a comparison to proposed steps of DreaM, **(C)** Steps for the deduplication and defined parameters related to read length and string match, separating deduped and duplicate reads, **(D)** Stats for the read deduplication on the CHM13 (CENP-A ChIP & CUT&RUN) after deduplication. **(E)** Panel showing the read base pair composition before and **(F)** after deduplication.

In these techniques, DNA-protein complexes are treated with cleaving enzymes or sonication to generate DNA fragments. These fragments are then sequenced to produce millions of short reads ranging from 75-151bp representing the genomic regions bound by the protein of interest^2,4–12^. The distribution and quantity of these reads across the genome provide insights into the location and strength of protein-DNA interactions, providing information on gene regulation^1,13^, chromatin structure^14– 18^, and epigenetic modifications^12,14,15,19–21^.

However, certain genomic regions, particularly those with repeated segments, presents significant challenges for these techniques. For instance; Centromeres are vital structures for accurate chromosome segregation during cell division, are composed of highly repetitive α-satellite DNA sequences (171bp) and unique epigenetic components^12,22–24^. Precise mapping and peak detection of protein-DNA interactions at centromeric regions or similar satellite such as human satellites pose persistent challenges due to the repetitive nature of sequences^12,16,23–29^ and the presence of polymerase chain reaction (PCR) duplicates within sequencing data^7,8,11,13,25,30–35^.

To overcome this challenge, we propose a pre-processing approach termed DreaM, which involves a FASTQ deduplication step prior to trimming in ChIP-Seq, CUT&RUN (**Fig 1B, C**). This early removal of duplicate reads reduces the impact of PCR duplicates, minimise multiple mapping and associated noise, which improves the accuracy of peak detection in these regions. Our comprehensive examination shows remarkable improvement offered by this approach compared to conventional pipeline, translating into highly significant peaks detected (q ≥0.00001 threshold) in narrow CDR region of centromere for both ChIP-seq and CUT&RUN studies.

## Results

### Pipeline design and quality control strategies in the DreaM method

To test the applicability of the approach, we utilized publicly available ChIP-Seq and CUT&RUN datasets targeting centromere protein A (CENP-A) from the CHM13 cell line, featuring sequence length of 151 bp. The workflow adopts initial deduplication step and follows a consistent series of steps, as depicted in **Fig (1B)**. To quantify the PCR duplicates in the tested dataset, total number of reads were counted in both deduplicated and duplicate groups which identified approximately 69.58% of the CENP-A enriched reads as PCR duplicates for ChIP-Seq and 45.91% for CUT&RUN (**Fig. 1D**). In contrast, the corresponding input data has only 6.0% of PCR duplicates, while IgG CUT&RUN contained 24.31% duplicates. (**Fig, D**).

To further analyse and represent the data, raw fastq reads were quantified by examining the percentage of bases (A, C, G, T, N) at each position before and after deduplication, presented as stacked plots (**Fig. 1E, F**). This analysis highlights changes in base representation following deduplication. We expected data with minimal duplicates to show a uniform base frequency similar to the input data (**Fig. 1E, F**). In contrast, data with PCR duplicates would lead to selective base enrichment contributed by similar reads, appearing as variable histograms for each individual base.

Given the multiple PCR amplification steps employed during library preparation, a high percentage of reads are expected to be PCR duplicates. This is likely due to the fact that centromeres are small regions within the genome compared to the total chromosome length. Consequently, the reads obtained through the ChIP-Seq and CUT&RUN protocols represent a very limited quantity, necessitating several cycles of PCR to prepare a sufficient sample for sequencing. These duplicates were successfully filtered out by the deduplication step as confirmed by FastQC (**Fig2A, B**).

**Figure 2:**
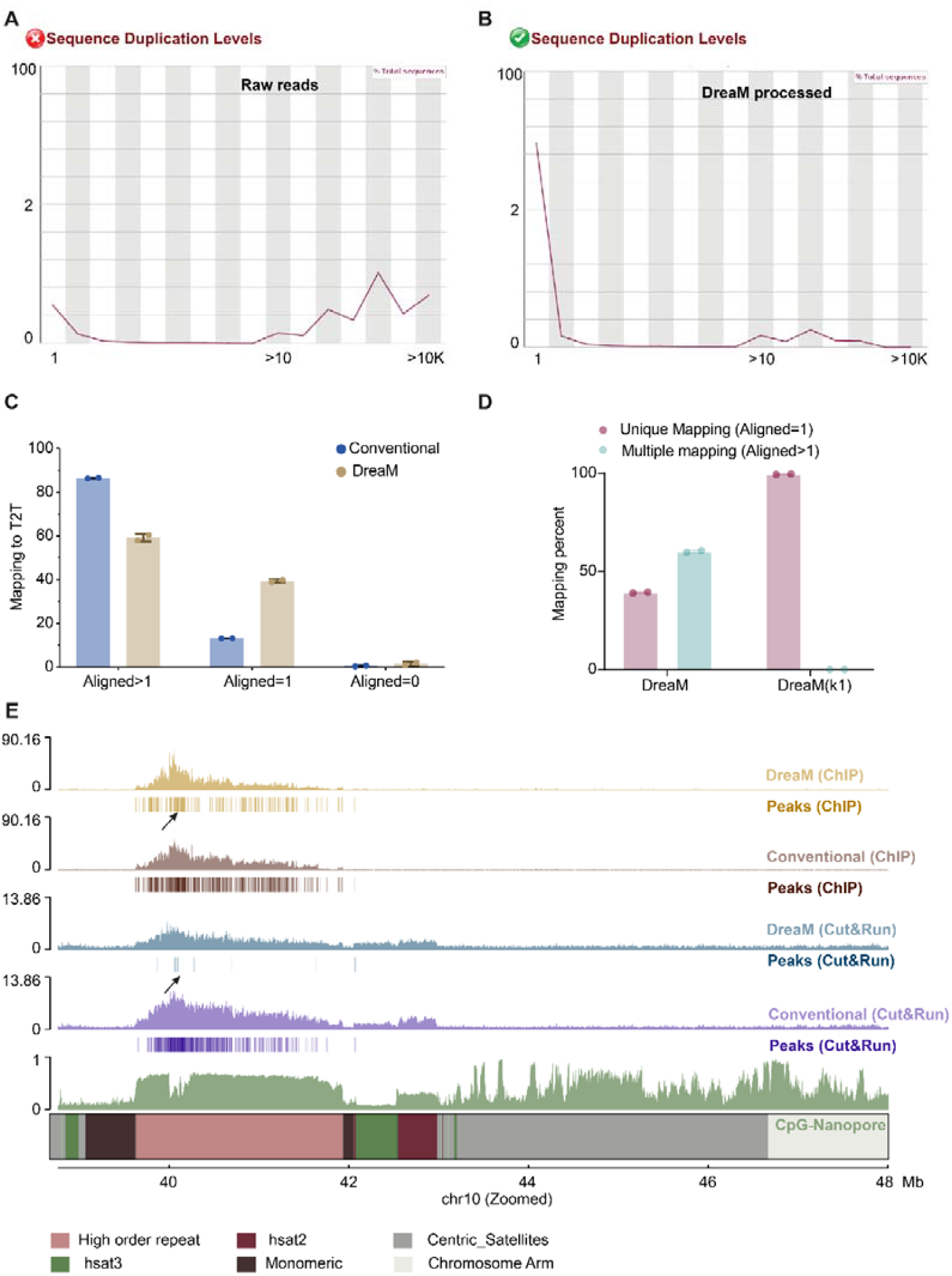
*FastQC* derived plots **(A)** before and **(B)** after deduplication showing the efficient removal PCR duplicate read stat passing the quality control, **(C)** Mapping statistics of raw reads and after DreaM’s processing shows decline in the multiple mapping (Aligned>1) and improvement in (Aligned=1, unique) upon deduplication. **(D)** utilizing the “k=1” parameter in Bowtie2 to retain reads with the best alignment score avoids multiple mapping completely. Panels Error bars represent the mean with SD among replicates. **(E)** ChromPlot generated by pyGenomeTracks, showing enhanced peak signal detection within the CDR (hypomethylated region) illustrated by the CpG nanopore data and detected peaks.

### DreaM Outperforms conventional method for Peak Detection

The DreaM pipeline outperforms conventional methods for peak detection, particularly in challenging regions like centromeres, where multiple mappings traditionally contribute to noise. Deduplicated reads were trimmed using *Trimmomatic* and mapped to the gapless T2T genome with *Bowtie2*, following established protocols^31,33,35–37^, achieving a 98% overall alignment rate across datasets. A detailed analysis revealed a 30% reduction in the proportion of reads mapping to multiple locations after deduplication, resulting in a higher proportion of unique alignments (**Fig 2C**). This pre-processing step effectively mitigated 30% of multiple mapping events, a major source of noise in conventional pipelines. Additionally, by controlling for multiple mappings (k=1), only the single best alignment was reported (**Fig 2D**).

Further, we used *MACS3* to call peaks and bamCoverage for computing overall coverage with CPM normalization, visualizing the results with pyGenometracks (**Fig 2E**). To highlight the improvement in CENP-A peak enrichment compared to conventional strategies, we compared the CENP-A coverage and peaks with CpG coverage from previously reported long-read sequencing data^9,29,36,38^. As anticipated, peaks are now more selectively enriched in hypomethylated regions of the centromere compared to distribution found in conventional pipeline in both ChIP and CUT&RUN datasets. Notably, we have employed a very stringent threshold of (q-value ≤0.00001) instead of 0.05 to detect peaks with high confidence, compared to conventional pipelines.

## Discussion

Efforts to dissect the complexity of the human genome demands ongoing improvements in sequencing technologies and bioinformatics algorithms to explore the functionality of genomic regions^30^. The initial release of the Human Genome Project in 2003 provided critical insights but focused primarily on coding regions, leaving 8% of centromeric/repetitive regions unexplored as “black box” gaps^10,30^. Recent advances such as Oxford Nanopore Technology (ONT)^6,39^ and PacBio^40^ facilitating advancement in technology for long-read sequencing aided to generate gapless reference genome Telomere-to-Telomere (T2T)^9,12,41^ assembly marking significant milestones by including previously inaccessible regions such as centromeric satellite arrays^23,28^ and segmental duplications^9,42–46^. This breakthrough allows for the targeted study of genomic variations by facilitating the mapping of short reads to previously difficult-to-explore regions such as centromeres, formerly disregarded as “blacklisted”^17,25,47–52^.

Despite the fundamental importance of centromeres in chromosome transmission during cell division, mapping and peak detection of these regions remains highly challenging due to repetitive sequences and PCR duplicates within short-read sequencing datasets. Hence, k-mer based approaches ^36,53^ are equally popular but produce similar results, as PCR duplicates may contribute to the enrichment of these k-mers originating from the same sequences, and do not offer a complete remedy for multiple mapping of k-mers at later stage, hence is used as a parallel confirmation.

We argue that the standard practice of removing PCR duplicates post-mapping may not be effective in avoiding issues such as multiple mappings and biases in specific genomic regions, potentially compromising peak calling accuracy^35,38,54,55^. To address this challenge, we propose implementing a FASTQ deduplication using the custom AWK algorithm (**Fig 1B, C**). This method effectively identifies PCR duplicates by evaluating two key criteria: the exact match in read length, and the identical string sequences (**Fig.1C**). The rationale for using these two parameters is that reads of exactly the same length and string match (base pairs) are more likely to be mapped to the same genomic coordinates, which in a standard pipeline is considered as PCR duplicates. We have shown that upon implementing the deduplication before attempting the mapping step, severely reduces the multiple mapping and aids in detecting high confidence peaks for centromere (**Fig 2E**).

### Advantages of DreaM

The shift towards long-read sequencing is inevitable due to its superior resolution, but the transition is gradual and driven by technological advancements. Significant cost reductions are needed to make it affordable for all. Until then, short-read sequencing will remain the gold standard, offering accurate, cost-effective solutions for large-scale projects, despite limitations in repeated regions that DreaM addresses to much extent.

Our approach offers several key advantages: it operates without virtual environments, simplifying workflows with a hassle-free command line interface; requires no updates, ensuring stable processing without compatibility issues; and processes large datasets efficiently, handling 65,658,018 header lines in a FASTQ file within 5-6 minutes using just 1 CPU and 8GB RAM, making it resource-efficient. Additionally, DreaM opens tempting doors and allows for the re-examination of existing data by applying to repeated regions in combination to T2T gapless reference genome to gain new insights, maximizing the value of previously generated data.

### Limitations

Our approach is expected to work on ChIP and CUT&RUN datasets provided the sequencing has been performed in depth and have read length sufficiently long as 150bp. We have tested this on GLOE-Seq or END-seq (data not shown). Unfortunately, these datasets are having more than 85% of duplicates and depth is not adequate leaving very less reads to be analyzed or mapped to genome. Also, this pipeline does not remove the optical duplicates that is present at minimal, this is addressed after mapping step only and is removed before peak calling.

## Methods

### Detection and deduplication of sequencing reads using custom AWK Script

Sequencing data generated from high-throughput platforms such as next-generation sequencing (NGS) often contain duplicate reads due to technical artifacts or PCR amplification biases. To address this issue and optimize downstream analyses, a custom AWK (Aho, Weinberger, and Kernighan)^56^ script was developed for detecting and deduplicating duplicate reads based on sequence content and length.

#### AWK Script Implementation

The AWK script utilizes a custom parsing strategy to process FASTQ files and identify duplicate reads based on their sequence content and length. The script executes as a standalone tool using the AWK interpreter, and the specific implementation details are outlined below:

#### Description of AWK Script Functionality

The AWK script processes input FASTQ files, where each record represents a sequence read and its associated quality scores. Key functionalities of the script include; (1) Parsing and processing of each record line by line, where sequences and quality scores are stored and read based on predefined conditions. (2) Identification of unique reads by checking sequence length and content, using a hash-based approach (seen array) to track previously encountered sequences. (3) Deduplication of reads into separate output files (_dedup.fastq and _duplicate.fastq) based on uniqueness criteria defined during processing. An average time taken for processing data with (65658018 fastq header lines) takes approx. (real 6m42.323s, user 5m36.603s, sys 0m21.097s) using 1 CPU with 8GB RAM as computed (time awk -f $script.awk /$file.fastq).

The script “DreaM.awk” can be executed by two ways. With command line (*for file in \**.*fastq; do awk -f* .*/DreaM*.*awk “$file”;done*), where each file in the folder with .fastq file extension will be processed and each read is handled separately as single-end and treated independent of each other for deduplication. In case of paired end, once processed as single end reads, both dedup files are compared to each other for match pairing based on headers and synced to get equal number of reads paired as forward and reverse file using script “Sync_DreaM.awk” which will produce output as $file_synced_R1 & $file_synced_R2.fastq. Commands can be executed as follows (*awk -f* .*/Sync_DreaM*.*awk “$file_R1_dedup*.*fastq” “$file_R2_dedup*.*fastq”*). These files can further be trimmed using the trimmers of choice with paired end mode.

### Analysis of ChIP-seq and CUT&RUN

Upon deduplication using the “DreaM.awk”, the reads were tested for quality controls (for file in *_dedup.fastq; do fastqc “$file” -o ./fastqc/;done) and trimmed using the *Trimmomatic*/*Trim Galore* in preferred mode (single/paired-end mode) and aligned using *bowtie2* (As single/paired end mode), keeping the minimum size to be 100bp after trimming (refer to GitHub for detailed codes for individual datasets), following which alignment was performed with the parameters (bowtie2--end-to-end-x [index_reference] [read1.fastq] [read2.fastq]). Resulting SAM file was converted to BAM using *samtools*. Further, BAM is sorted, fixmate was applied and marked for any optical duplicates present, not addressed during deduplication. Later filtered for Flag 2308 (by using -F 2308 in *samtools*, that ensures to remove all reads except primary mapped reads) retains in the resulting file. This BAM file is used for further peak calling. All the analysis in this manuscript followed the standard pipeline and compared to DreaM approach. The versions of the algorithms applied are listed in the resource table and necessary codes are available on GitHub depository (https://github.com/santoshbiowarrior333/DreaM)

### Peak calling and visualization

The resulting files obtained from the (−F 2308) filtration step that is uniquely mapped reads, was used to call for peaks using *MACS3* with parameter specifying narrow/broad peaks. Significantly enriched (removing FLAG 2308 and retaining peaks with q-value ≤0.00001, parameters -f BAMPE -B -g 3.03e9 -q 0.00001)^57^ was computed resulting list of the peaks. Relative bigwig files were generated with deepTools and were plotted using the pyGenomeTracks. The versions of the algorithms applied are listed in the resource table.

## Data availability

The data analysis in this manuscript is performed on the published datasets from public depository (SRA), all the accession numbers are available in the resource table along with the cell line information.

## Code

All the codes used in the above manuscript is available on github for reference (https://github.com/santoshbiowarrior333/DreaM).

## Acknowledgement

Authors are thankful to Central Oxford Structural Molecular Imaging Centre (COSMIC), Kavli Institute for Nanodiscovery at University of oxford for providing the computational facility and support. We performed all the data analysis using the facility clusters. This work was supported by the Medical Research Council (MR/W017601, to F.E.) and the Edward Penley Abraham Research Fund.

## Resource table

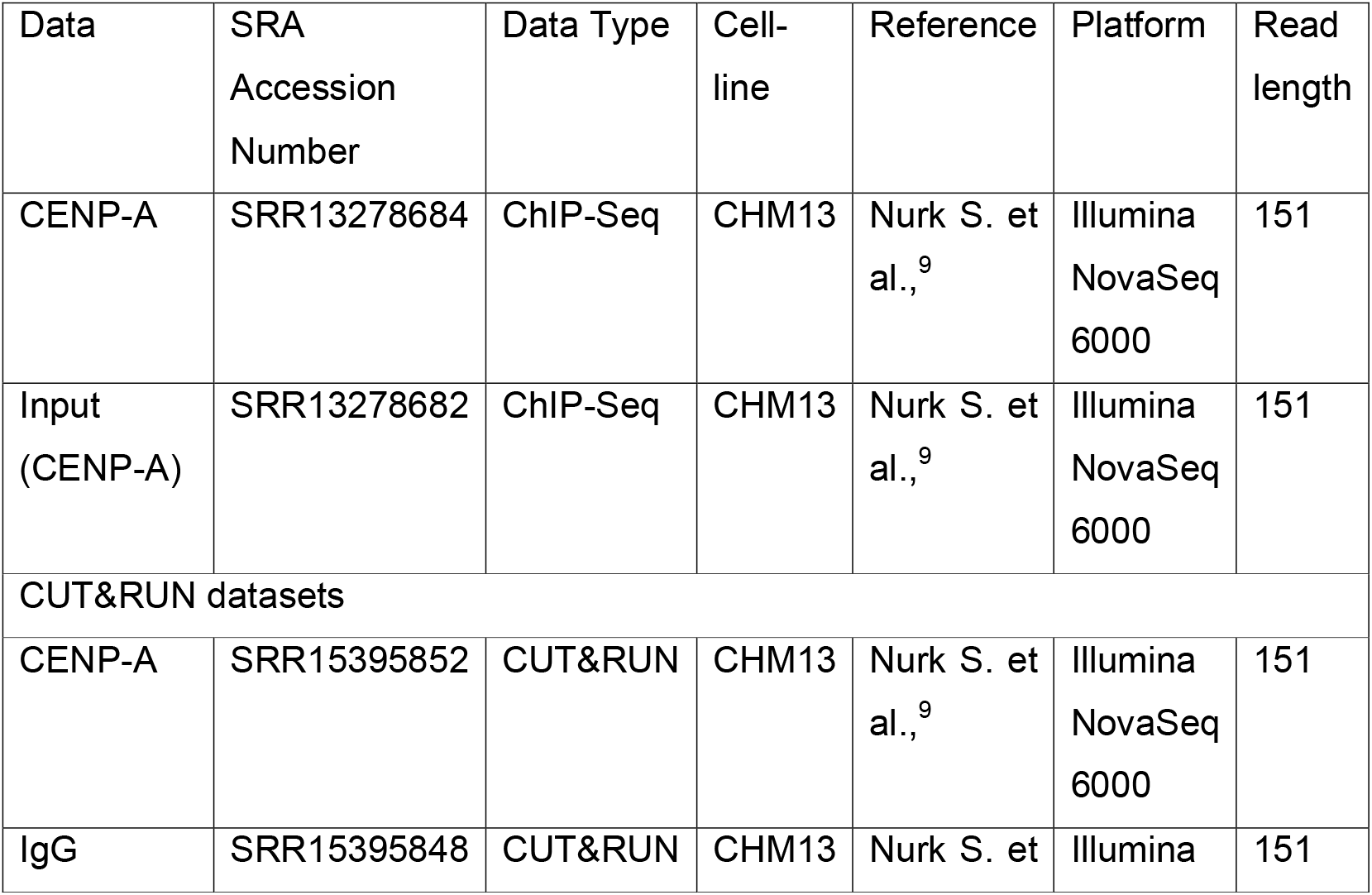
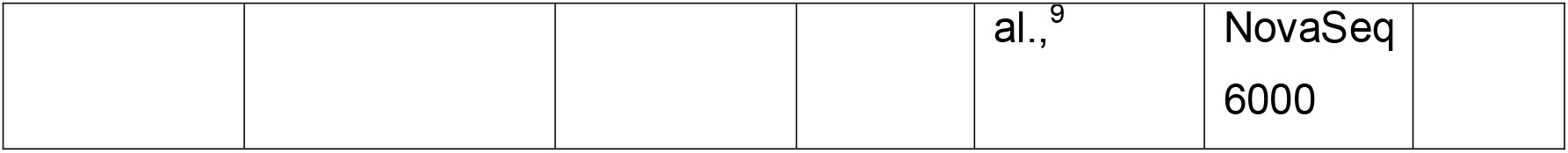

## Software and their versions

| Software and algorithms | References | RRID |
| --- | --- | --- |
| Trimmomatic v0.39 | Bolger et al<sup>58</sup> | RRID:SCR_011848 |
| FASTQC v0.12 | Babraham<br>Bioinformatics<sup>59</sup> | RRID:SCR_014583 |
| Bowtie2 | Langmead, B<sup>60</sup> | RRID:SCR_016368 |
| SAMtools v1.18 | Li et al<sup>61</sup> | RRID:SCR_002105 |
| deepTools v3.5.3 | Ramírez et al.<sup>62</sup> | RRID:SCR_016366 |
| GraphPAD Prism v10 | NA | RRID:SCR_002798 |
| Python v3.11.9 | Van Rossum and Drake | RRID:SCR_008394 |
| Matplotlib v3.8.4 | Hunter<sup>63</sup> | RRID:SCR_008624 |
| NumPy v1.26.4 | Charles R. Harris<sup>64</sup> | RRID:SCR_00863 |
| Pandas v2.2.2 | McKinney<sup>65</sup> | RRID:SCR_018214 |
| pyGenomeTracks v3.6 | Lucille Lopez-Delisle<sup>66</sup> |  |
| Seaborn v0.13.2 | Michael L. Waskom<sup>67</sup> | RRID:SCR_018132 |
| SEQTK | Wei Shen<sup>68</sup> | RRID:SCR_018927 |
| MACS3 | Yong Zhang et al<sup>55</sup> | RRID:SCR_013291 |

